# Recognition of a Tandem Lesion by DNA Glycosylases Explored Combining Molecular Dynamics and Machine Learning

**DOI:** 10.1101/2021.01.06.425536

**Authors:** Emmanuelle Bignon, Natacha Gillet, Chen-Hui Chan, Tao Jiang, Antonio Monari, Elise Dumont

## Abstract

The combination of several closely spaced DNA lesions, which can be induced by a single radical hit, constitutes a hallmark in the DNA damage landscape and radiation chemistry. The occurrence of such tandem base lesions give rise to a strong coupling with the double helix degrees of freedom and induce important structural deformations, in contrast to DNA strands containing a single oxidized nucleobase. Although such complex lesions are known to be refractory to repair by DNA glycosylases, there is still a lack of structural evidence to rationalize these phenomena. In this contribution, we explore, by numerical modeling and molecular simulations, the behavior of the bacterial glycosylase responsible for base excision repair (MutM), specialized in excising oxidatively-damaged defects such as 7,8-dihydro-8-oxoguanine (8-oxoG). The difference in lesion recognition between a simple damage and a tandem lesions featuring an additional abasic site is assessed at atomistic resolution owing to microsecond molecular dynamics simulation and machine learning postprocessing, allowing to extensively pinpoint crucial differences in the interaction patterns of the damaged bases. This work advocates for the use of such high throughput numerical simulations for exploring the complex combinatorial chemistry of tandem DNA lesions repair and more generally multiple damaged sites of the utmost significance in radiation chemistry.

## Introduction

The chemical stability of DNA components is fundamental to maintain the genome stability, hence preventing unwanted mutations or cell death. Indeed, the accumulation of DNA lesions has been recognized as one of the principal causes of cancer development^1^. Although DNA maximizes its stability through its helical structure, its constituting nucleic acids are constantly exposed to damaging agents, either endogenous or exogenous, that inevitably lead to the production of lesions. Among the different sources of DNA lesions, we may briefly remind oxidative agents, such as free radicals or reactive oxygen species (ROS), UV light, and ionizing radiations. As a consequence, specific and highly efficient repair machineries exist that are able to recognize the presence of lesions in the genome and remove them to reinstate undamaged DNA strands^2,3^. Specific DNA repair pathways may depend on the organisms, and are also related to the kind of lesions, for instance for localized oxidatively-induced damages the base excision repair (BER) pathway is preferred^4,5^, while for more extended and bulky lesions, such as base dimerization, the nucleotide excision repair (NER) mechanism is favored.

Yet this sophisticated repair mechanism has been reported to be strongly impaired when not only one but two adjacent DNA lesions are located on the same strand, the so-called tandem lesions. The formation of tandem lesions can derive from a single radical hit, and their biological impact is now well established. While their formation mechanism has been delineated^6^, the reasons underlying their resistance to repair are more elusive and should be analyzed taking firmly into account specific structural modification. The most common oxidative tandem lesions feature two adjacent oxidized nucleobases. In the following we will specifically consider 8-oxoguanine (8-oxoG) and an abasic apurinic/apyrimidinic site (Ap), as shown in Figure 1-C. This arrangement is particularly relevant also because Ap are also the most common outcome of ionizing radiations after excision of an entire nucleobase.^7^. Interestingly, Ap sites also represent key intermediates of the BER machinery and result from the action of DNA glycosylases before being further processed and removed by endonucleases. The presence of a tandem lesion, or more generally multiple damaged sites (MDS), that are the hallmarks of radiation chemistry, induces strong coupling between the lesions that in turn is translated into important structural deformations of the nucleic acid as compared to its ideal structure, i.e. either undamaged strand or sequences containing an isolated lesion. The unusual structural deformations induced by tandem lesions or MDS may also well justify their globally lower repair rate as compared to other lesions^8–11^.

**Figure 1.**
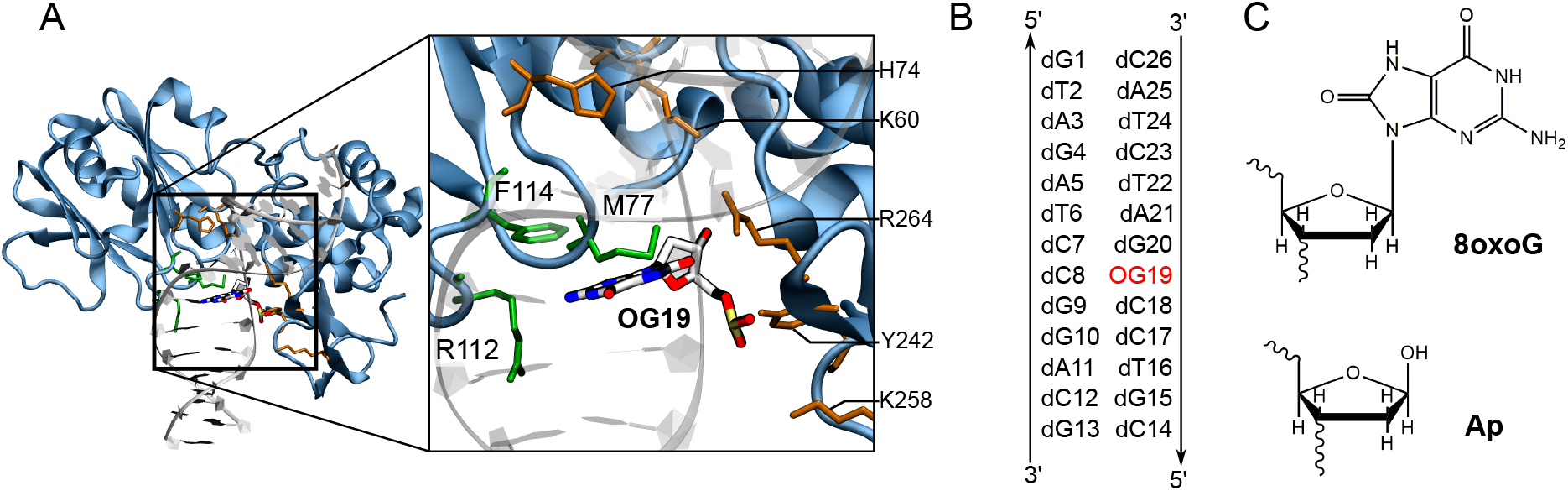
(**A**) Cartoon representation of the bacterial MutM in interaction with a 13-bp double stranded DNA helix harboring 8-oxoguanine (OG19) as the 19th nucleobase — PDB ID code 3GO8^21^. The magnified section highlights the position of the catalytic triad (M77, R112, and F114 in green) and the residues interacting with the DNA backbone (in orange) around the damage. (**B**) Sequence of the 13-bp oligonucleotide, showing the position of the 8-oxoguanine (in red). In simulations with tandem lesions, dG20 is mutated *in silico* into an abasic site (Ap). (**C**) Chemical structure of the 8-oxoguanine and the abasic site lesions.

To cope with their frequency, canonical DNA lesions benefit from a most efficient repair. For instance, 8-oxoG, which is well-known to mismatch with adenine and hence is potentially mutagenic^12^, is repaired by formamidopyrimidine DNA glycosylase, an enzyme that is referred to as Fpg in eukaryotes, while its bacterial counterpart is called MutM. The latter recognizes the presence of 8-oxoG in the genome and specifically binds at the damaged site^13^. Many studies have contributed to dissect the mode of action of MutM/Fpg in presence of a single 8-oxoG in particular concerning the recognition of the lesion^14–16^. Fpg^17,18^ has been shown to recognize 8-oxoG among other oxidatively-induced lesions and to subsequently proceed to its extrusion initiating the base excision process^4,5^. The mechanisms of recognition^19^ and extrusion^17,20,21^ of 8-oxoG have been scrutinized through a series of techniques, including molecular modeling and simulations, and are now relatively well characterized. Recently, Simmerling et al.^22^, while recognizing the role of the damaged base flipping in favoring its recognition, have also pinpointed the existence of preliminary recognition steps correlating with the rapid sliding of Fpg along the DNA strand that is incompatible with a recognition mechanism based on the systematic flip of all the bases. In addition, the same authors have also identified that 8-oxoG flips preferably through the major groove. The free energy required for the extrusion of 8-oxoG in extrahelical position has also been estimated by La Rosa and Zacharias^16^, also taking into account the contributions due to the DNA global bending and twisting. A most important feature of MutM/Fpg efficiency has been traced back to the crucial M77, R112, and F114 amino acids triad. Indeed, it permits to disrupt 8-oxoG interactions within the DNA helix by intercalating above the 8-oxoG position, thereby facilitating its extrusion towards the active site. Besides, other several important MutM/Fpg residues (K60, H74, Y242, K258, and R264) are known to stabilize the DNA helix by interacting with its backbone^20^.

On the other hand, several studies have addressed the behavior of tandem-containing oligonucleotides, either from a biochemical and repair perspective^9^ or from a structural point of view^23^, also relying on molecular modeling and simulations^11,24,25^. Globally, the different approaches agree in pointing out a strong effect of closely spaced lesions in modifying the structure and dynamics of the oligonucleotide. In addition, strong sequence effects, depending both on the relative position of the cluster lesions and on the nearby undamaged bases contribute to the complexity of the global landscape. The interaction of MDS-containing oligonucleotides with repair enzymes and in particular both *E. Coli* and human endonucleases^10,26^ has been reported. The perturbations exerted by the secondary lesion on the protein/DNA contact regions, and the consequent decrease in its repair efficiency, as observed for some particular tandem lesions have also been highlighted. However, no analysis of the structural behavior of Fpg and MutM in presence of tandem DNA lesions has been reported, despite the relevance that such lesions may assume in conditions of strong oxidative stress or ionizing radiations.

In this work, we take advantage of the existing knowledge of 8-oxoG recognition by MutM to investigate the structural and dynamic impact of the presence of a second, adjacent lesion, namely an Ap site. Relying on all-atom, explicit-solvent molecular dynamics^10,27^ we simulate the structural and dynamical behavior of a MutM:DNA complex. We consider both the situation in which only a single lesion (8-oxoG) is present and compare it to the one in which the adjacent guanine base dG20, present in the X-ray structure (PDB ID 3GO8), has been *in silico* mutated to an Ap site - see Figure 1-B. We clearly show that the presence of the tandem lesion induces important structural deformations to the DNA that significantly perturb the protein/nucleic acid interaction pattern, hence being susceptible to alter the 8-oxoG extrusion. We extensively describe the changes in the 8-oxoG lesion structural signature upon the presence of an adjacent Ap site and the perturbation of the interaction network with MutM (Fpg), which contribute to ultimately diminish the recognition efficiency.

## Results

We report the structural and dynamical properties of the bacterial MutM interacting with a 13-bp DNA sequence harboring either a single 8-oxoG at the position 19 (as found in the crystal structure PDB ID 3GO8) or 8-oxoG coupled with an Ap site at position 20, along two replicas reaching 1*μ*s MD simulation time each. The numbering of the nucleic acids used hereafter corresponds to the one in Figure 1-B; the numbering of MutM residues refers to the crystallographic structure (PDB ID 3GO8).

### Tandem lesions impact the interaction network around 8-oxoG

The interaction network as found in the MutM:DNA crystal containing a single 8-oxoG lesion is conserved stable along our MD simulations. A most important structural feature in MutM is its intercalation triad, consisting of the M77, R112 and F114 residues. Those three amino acids are located around the 8-oxoG in the minor groove, weakening the stabilizing interactions of the lesion within the double-helix to facilitate its extrusion. R112 interacts with the complementary dC7, while M77 and F114 intercalate directly above 8-oxoG and disrupt the stable *π*-stacking with the adjacent base-pair – see Figure 2-A. These interactions are persistent along the entire MD simulations of the singly-damaged system. F114 is involved in *π*-staking with dG20 during 91.8% of the time series, with the distance between heavy atoms of their aromatic rings averaging at 5.35±0.5 Å. The rest of the time, F114 stacks transiently with dC7 facing dG20 and their aromatic rings maintain a distance of 6.7±0.5 Å along the simulations – see Figure 3. M77 intercalates between OG19 and dG20, as its terminal methyl group remains at 4.8±0.4 Å of the N9 atom involved in the N-glycosidic bond, and is ideally positioned to act on OG19 desoxyribose moiety to drag it outwards^20^.

**Figure 2.**
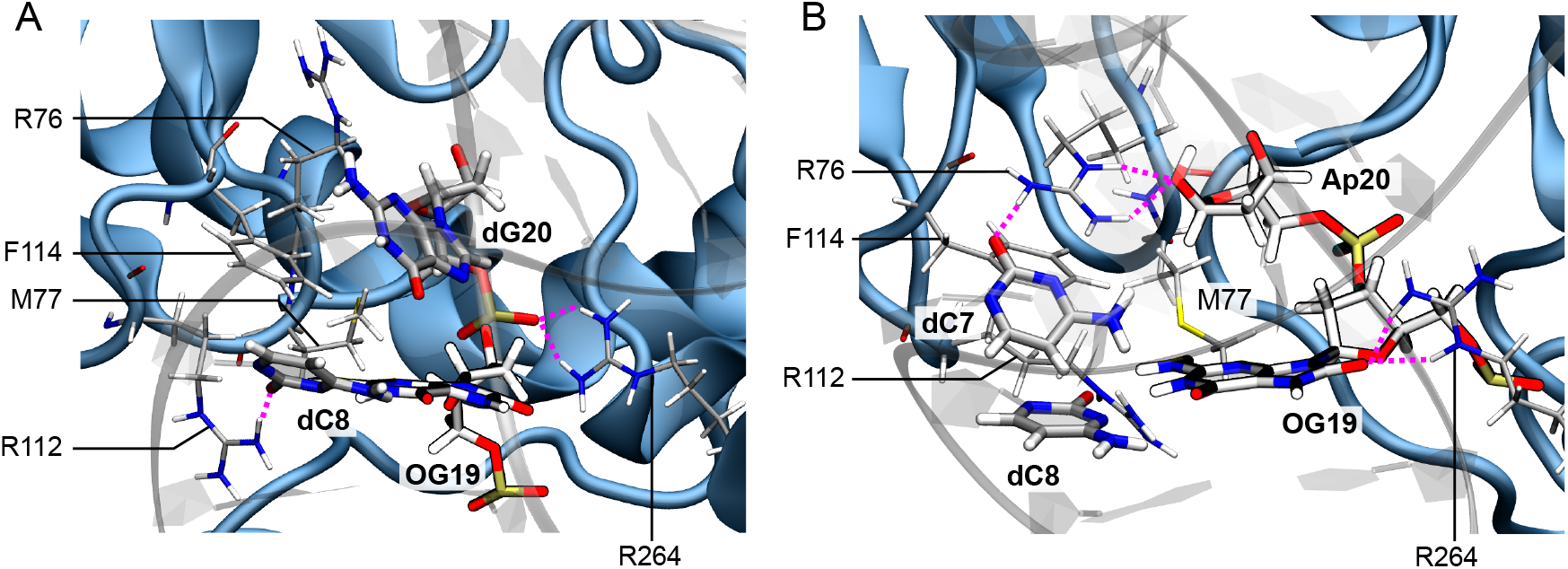
Cartoon representation of MutM interacting with the DNA helix harboring a single 8-oxoG lesion (OG19, A) or tandem lesions 8-oxoG + Ap (OG19 and Ap20, B). H-bonds are depicted as dashes pink lines and the DNA structure is rendered transparent for sake of clarity. Upon multiple lesions, the interaction pattern around 8-oxoG (OG19) is perturbed. The intercalation triad M77/R112/F114 is shifted down by R76 which comes to interact between Ap20 and the facing dC7, preventing M77 ad F114 intercalation above 8-oxoG. R264, normally interacting with the DNA backbone phosphates between positions 19 and 20, is now involved in hydrogen bonding with OG19 carbonyl.

**Figure 3.**
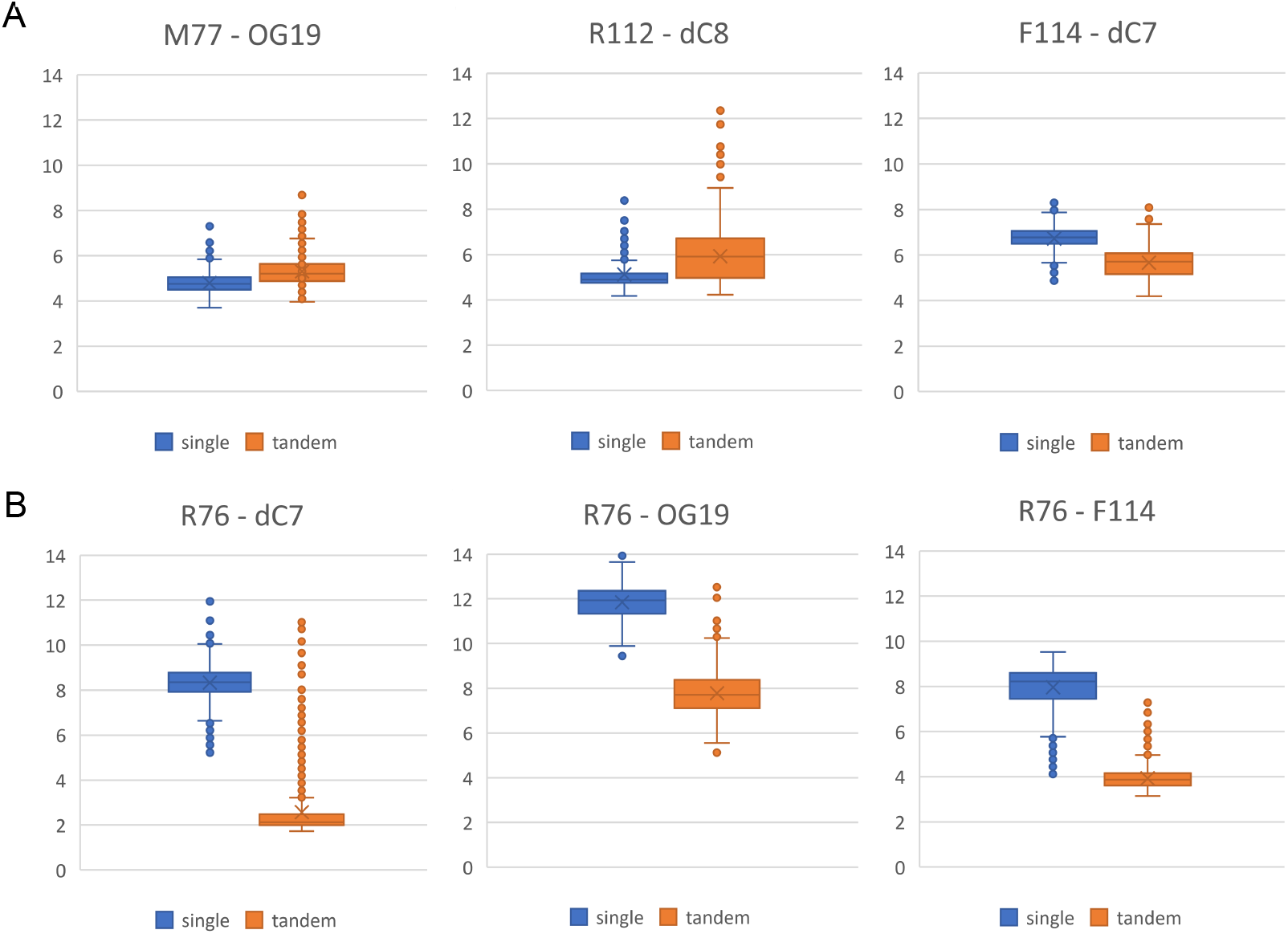
Distribution of relevant distances involving the intercalation triad (A) and R76 (B) upon a single 8-oxoG mutation (single) or Ap + 8-oxoG lesions (tandem). The presence of the Ap site at position 20 makes M77 and R112 move away from 8-oxoG and the facing dC8. F114 makes *π*-stacking with dC7 because the nucleobase at position 20 is now absent. R76 comes closer to 8-oxoG and intercalates in the gap left by the abasic site, with formation of very stable H-bonds bridging Ap20 and the facing dC7. It also interacts with F114, preventing it to stack within the double-helix.

R112 side chain amino groups form H-bonds with the nitrogen and carbonyl of dC8 over 76.8% of the simulation time, the distance between these two atom groups being of 5.1±0.6 Å. Several other amino acids have been identified to stabilize the MutM:DNA complex by interacting with the negatively-charged phosphate groups of the backbone namely K60, H74, Y242, K258, and R264. These interactions are stable in our simulations and the highly conserved R264 forms strong H-bonds between OG19 and dG20 phosphate groups - as shown in Figure S1. Noteworthy, R264 is known to play a role in 8-oxoG extrusion^22^. The structural behavior of MutM:DNA(8-oxoG) observed here corroborates the hypothesis of a highly dynamic system, whose functional flexibility is known to be central to ensure its biological role through the recognition and extrusion of 8-oxoG^28,29^.

The dG20 → Ap20 mutation induces a clear perturbation of this well-characterized interaction network. A first consequence is the perturbation of the dynamics of the intercalation triad. The presence of the abasic site involves, in the first 100 ns of simulation, a rapid reorientation of R76 situated just above the intercalation triad. R76 side chain turns towards the damaged site, and is found closer to OG19, at 7.8±0.8 Å vs. 11.8±0.7 Å observed in the singly 8-oxoG-containing duplex. R76 does not interact directly with OG19 but rather positions itself in the gap between the Ap site and the facing dC7, bridging the two residues through stable H-bonds as reported in Figure 2-B. The distance between the R76 guanidinium nitrogens and the dC7/Ap20 H-bond acceptor atoms lies at 2.6±1.1 Å and 3.0±1.8 Å, respectively, in the tandem-damaged MutM:DNA complex. Comparatively, the dC7-R76 distance is of 8.3±0.7 Å in the singly damaged system – see Figure 3-B. The reorientation of R76 reshapes the canonical interaction network of the intercalation triad, which is globally shifted downwards the duplex. F114 is pushed away from position 20 and comes closer to the opposite strand, the dC7-F114 distance drops to 5.7±0.6 Å, although its strong cation-*π* interaction with R76 avoids direct stacking with dC7. Additionally, the distance between the R76 guanidinium extremity and the F114 aromatic ring is of 3.9±0.5 Å vs. 8.0±0.9 Å in the singly-damaged system, while the interaction of R112 with the estranged dC8 is destabilized. In presence of tandem lesions, R112 lies further from dC8 (5.9±1.0 Å) than what is observed for the singly-damaged complex (5.1±0.6 Å). The intercalation of M77 is prevented in presence of the tandem lesion since its terminal methyl group rotates away from OG19:N9 (5.3±0.6 Å), while the interaction with M77 corresponds to a more rigid binding mode, with the formation of a H-bond between the sulfur atom and one hydrogen of OG19. The corresponding distance is reduced to 2.7±0.7 Å vs. 3.5±0.8 Å with the singly damaged (OG19) duplex.

Interactions between the DNA backbone and MutM tend to be more rigid upon tandem damages than in the singly damaged duplex. The H-bond between K60 and the phosphate at position 20 is stronger as witnessed by the −NH_3_^+^... P distance that is reduced to 5.3±1.1 Å vs. 8.2±2.6 Å for the singly-damaged system, as well as the interaction of H24 with dA21 (NH - P distance of 4.3±0.6 Å vs. 5.4±2.0 Å with the isolated 8-oxoG) and Y242 H-bond with OG19 (OH - P distance of 5.1±1.7 Å vs 7.1±2.3 Å). However, the interactions of R264 with the DNA helix is strongly perturbed: in the singly-damaged system, R264 forms stable H-bonds with OG19 and dG20 phosphates (CZ - P distance of 4.9±1.9 Å and 4.9±1.3 Å, respectively) which are disrupted upon the presence of the additional Ap site (CZ - P distance of 8.1±3.2 Å and 8.8±2.4 Å, respectively). The R264 position experiences important fluctuations in the tandem-damaged complex, and can form stable H-bond with OG19:O8 - see Figure 2-B. the R264:CZ - OG16:O8 distance is below 4 Å for 42% of the simulation time in the tandem-damaged complex, while in the singly-damaged complex such short distance amounts to 11% only - see Figure S2.

This first local analysis suggests that the singly-*vs.* tandem-damaged 13-bp duplex present different interaction patterns, with non trivial changes in the binding mode and its dynamics. In order to probe more extensively the structural and dynamic consequences of dG20 → Ap20 substitution, we have relied on a recently-proposed machine-learning protocol^30^ to identify other residues possibly implied in the recognition mechanism.

### Systematic assessment of interacting residues through machine-learning protocol

In order to probe the residues that exhibit important interactions with the DNA duplex, a machine-learning protocol based on the multilayer perceptrons (MLP) classifier was set up. The latter allows to generate a “footprint” of the residues that are particularly involved in MutM:DNA bonding – see Figure 4. A score function, in the following referred to as ‘importance’, is attributed to each residue: the higher the score, the higher the contribution to the MutM:DNA complex stabilization. Using a threshold of 0.04 of importance, 47 and 61 residues out of 273 single out in the singly and tandem-damaged system, respectively.

**Figure 4.**
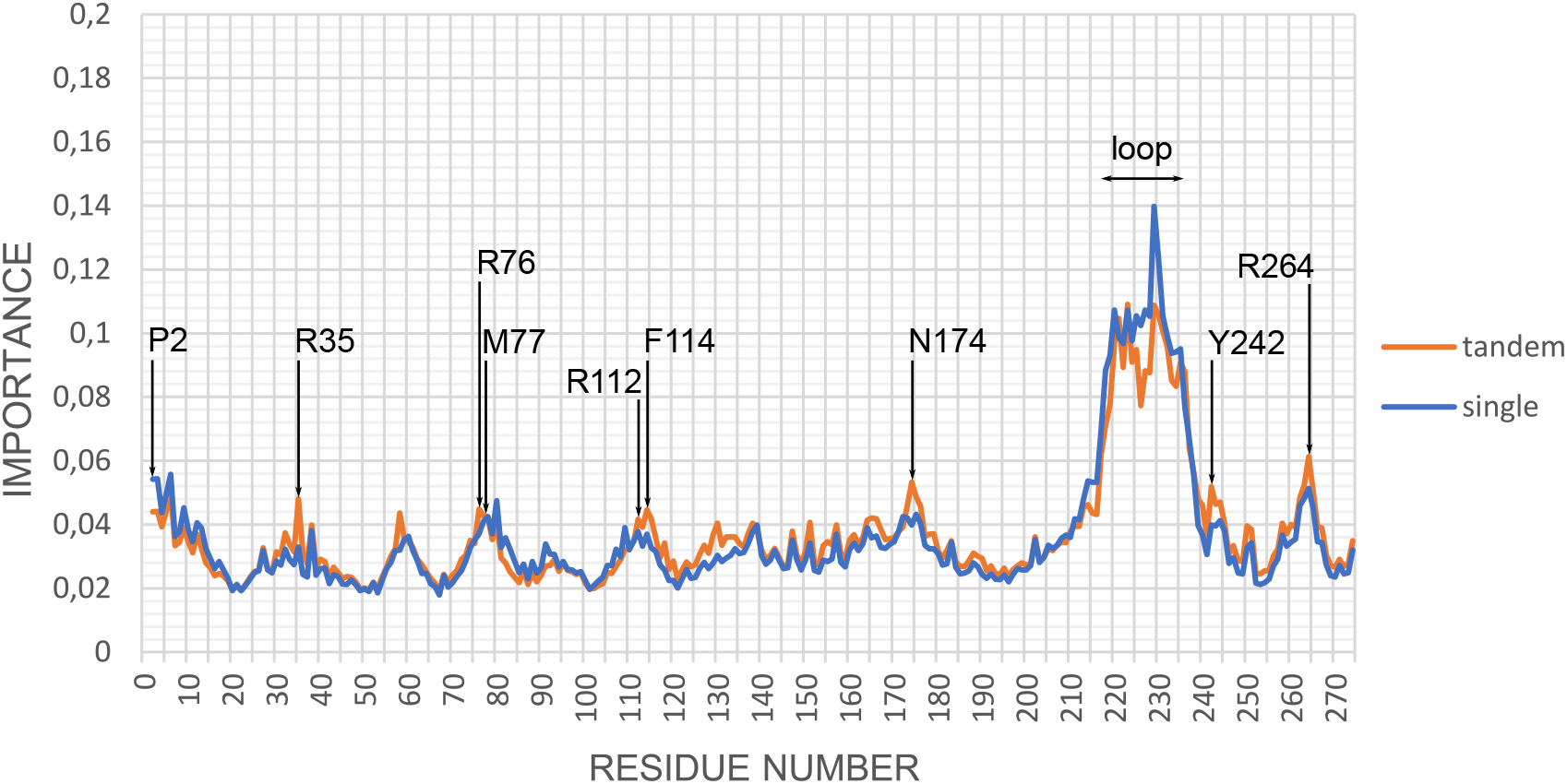
Importance of the contribution of residues to the MutM:DNA complex bonding for the singly-damaged (blue) and the tandem-damaged (orange) systems. The threshold value above which the importance of the residue for the stabilization of the complex is considered as significant is 0.04. Some of the key-residues as well as the flexible loop region are pinpointed by the arrows. Contributions of amino acids to the bonding are mostly higher upon 8-oxoG/Ap combination, suggesting a more rigid complex upon multiple damage sites than with an isolated 8-oxoG.

The three residues of the intercalation triad (i.e. M77, R112 and F114) show a slightly higher contribution in the tandem-(0.043, 0.042, 0.045) than in the singly-damaged system (0.041, 0.038, 0.037). R76 and Q78, adjacent to M77 in the MutM sequence, also present high values. As highlighted by the visual inspection of our MD trajectories, in the tandem-damaged system, R76 flips towards the lesion site to compensate for the nucleobase removal at position 20 by bridging Ap20 to the facing dC7 through strong H-bonds. The importance score for R76 is 0.045 with tandem lesions vs 0.037 with isolated 8-oxoG, corroborating the significant role of this residue in MutM:DNA binding upon the presence of Ap20, in line with the newly-formed and very stable H-bonds with the lesion site. Q78 importance is higher than the threshold in both tandem-(0.042) and singly-(0.043) damaged systems. This residue interacts with R112, contributing to the H-bonds network in the vicinity of the lesion. Adjacent to F114, G115 importance in the stabilization of the complex is also enhanced upon dG20 → Ap20 mutation (0.042 in tandem vs. 0.033 with isolated 8-oxoG). Additional visual inspection of the MD trajectories reveals that G115 forms a strong H-bond with R76, helping in maintaining the latter intercalated between Ap20 and the facing dC7.

Over the five key residues reported to anchor the phosphate DNA backbone (K60, H74, Y242, K258, and R264^31^), only the closest to OG19 are associated with importance scores above the threshold of 0.04: Y242 (0.040 and 0.052 for the singly- and tandem-damaged system), K258 (0.037 and 0.040) and R264 (0.051 and 0.061). The G173 and Y176 residues also contribute to the H-bonding with the DNA backbone. N174 shows high importance values, 0.040 and 0.053 for singly- and tandem-damaged systems, this is due either to its interaction with the damaged site backbone or through indirect coupling with R264, as previously described in the literature^17^. I173 is involved in hydrophobic interactions with Y242 that in turn interacts with OG19 backbone. Among other residues whose contribution is above the threshold, R35 forms H-bonds with either dC8 or dC23 backbone, P130 and M166 interact with dT22:P, D165 and R150 maintain the 5’-terminus backbone of the DNA strand 1 (in the dG1 and dT2 surroundings), L164 stabilizes the position of the key-residue R264, while G263 and G265 form H-bonds with R264 or directly with the DNA backbone, and K258 interacts with the dC17 phosphate.

Globally the MLP analysis clearly reveals that the protein residues comprised between the position 210 and 237 exhibit the highest values of importance. They correspond mostly to a large, flexible loop, comprising the residues 221–234 at the C-terminus that is prone to disorder, but also known to have an implication for DNA recognition despite being spatially far from the double-helix^21^. Amino acids at the N-terminus also show significant contributions to the MutM:DNA bonding. The proline located at the very end of the N-terminal region has an important catalytic role since it reacts with the C1’ atom of the deoxyribose sugar moeity of the 8-oxoguanine to form a Schiff base, and hence it induces the cleavage of the N-glycosidic bond which constitutes the first step of the repair process. Adjacent to P2, the vicinal E3 is also known to play a role in MutM catalytic efficiency. Interestingly, the contribution of these two residues to the MutM:DNA stability decreases from 0.054 in the singly-damaged to 0.044 for the tandem-damaged complex, hence corroborating a subtle reduction of the excision efficiency. Other residues of the N-terminus (L4, P5, E6) also show a drop in their contribution upon dG20 *→* Ap20 mutation.

The residues which single out in this MLP analysis match very well with the ones evidenced by previous works on MutM and Fpg^14,15,17,19,20,28^. Our machine-learning post-processing allows to disentangle a complex interaction pathway, which is already well-established for 8-oxoG-containing DNA^29^ but perturbed upon the presence of tandem lesions as revealed by the present simulations. It allows to generate an exhaustive map of residues showing importance for the protein-DNA interactions, beyond the simple visual investigation based on the data from the literature. Noteworthy, the nucleic acid importance score in the MutM:DNA bonding is enhanced upon the presence of Ap20, denoting again a more constrained oligonucleotide - see Figure S3.

### Mechanical and dynamic properties of the DNA strand

In order to assess the mechanical and dynamic properties of the DNA strand, the MD trajectories were post-processed with the Curves+ program^32^ to evaluate the structural parameters of the double helix.

The first signature of the B-helix is often the bend angle, which reaches typical values around 51±11° upon interaction with MutM for the singly-damaged oligonucleotide. Such extreme values for bending are typical^33,34^ and necessary to facilitate the extrusion of the lesion towards the enzyme active site. The presence of the Ap site at position 20 is not sufficient to perturb the global bending of the 13-bp oligonucleotide (49±12^°^), but rather induces local deformations.

Structural parameters of the dC8-OG19 basepair are particularly impacted, with values lower for the tandem- than for the singly-damaged system - see Table 1. Importantly, the backbone parameters ‘Bend’, ‘Tip’ and ‘Inclination’ are lower when the Ap site is present at position 20, denoting a straighter portion of DNA helix than what is normally found in the canonical single-damaged MutM:DNA complex - see Figure 5 and Figure S4. The values monitored for these parameters are of 8.6±1.6°, 14.0±5.8°, and 19.5±4.3°, respectively in the singly-damaged system, vs 7.6±1.9°, 8.1±6.1°, and 18.3±7.0° in the tandem-damaged complex. Besides, several intra base-pair structural parameters are also found closer to the canonical B-DNA for the dC8-OG19 base-pair. Especially, the ‘Buckle’ and ‘Propeller’ drop from −16.1±8.4° to −8.8±10.7° and from 5.6±7.5° to 0.0±9.3°, respectively upon dG2 → Ap20 mutation. Other parameters (Opening, Shear, Stretch, and Stagger) show less significant deviations - see Table 1 and Figure S5.

**Table 1.** Averaged values of the dC8-OG19 base-pair structural parameters, for the single 8-oxoG (Single, left) and the tandem 8-oxoG+Ap (Tandem, right).

| dC8-OG19 parameter | Single | Tandem |
| --- | --- | --- |
| Local bending ($^\circ$) | $8.6 \pm 1.6$ | $7.6 \pm 1.9$ |
| Tip ($^\circ$) | $14.0 \pm 5.8$ | $8.1 \pm 6.1$ |
| Inclination ($^\circ$) | $19.5 \pm 4.3$ | $18.3 \pm 7.0$ |
| Buckle ($^\circ$) | $-16.1 \pm 8.4$ | $-8.8 \pm 10.7$ |
| Propel ($^\circ$) | $5.6 \pm 7.5$ | $0.0 \pm 9.3$ |
| Opening ($^\circ$) | $0.1 \pm 3.9$ | $1.0 \pm 3.6$ |
| Shear ($\text{\AA}$) | $-0.14 \pm 0.33$ | $-0.21 \pm 0.39$ |
| Stretch ($\text{\AA}$) | $0.05 \pm 0.12$ | $0.02 \pm 0.12$ |
| Stagger ($\text{\AA}$) | $0.54 \pm 0.33$ | $0.49 \pm 0.43$ |

**Figure 5.**
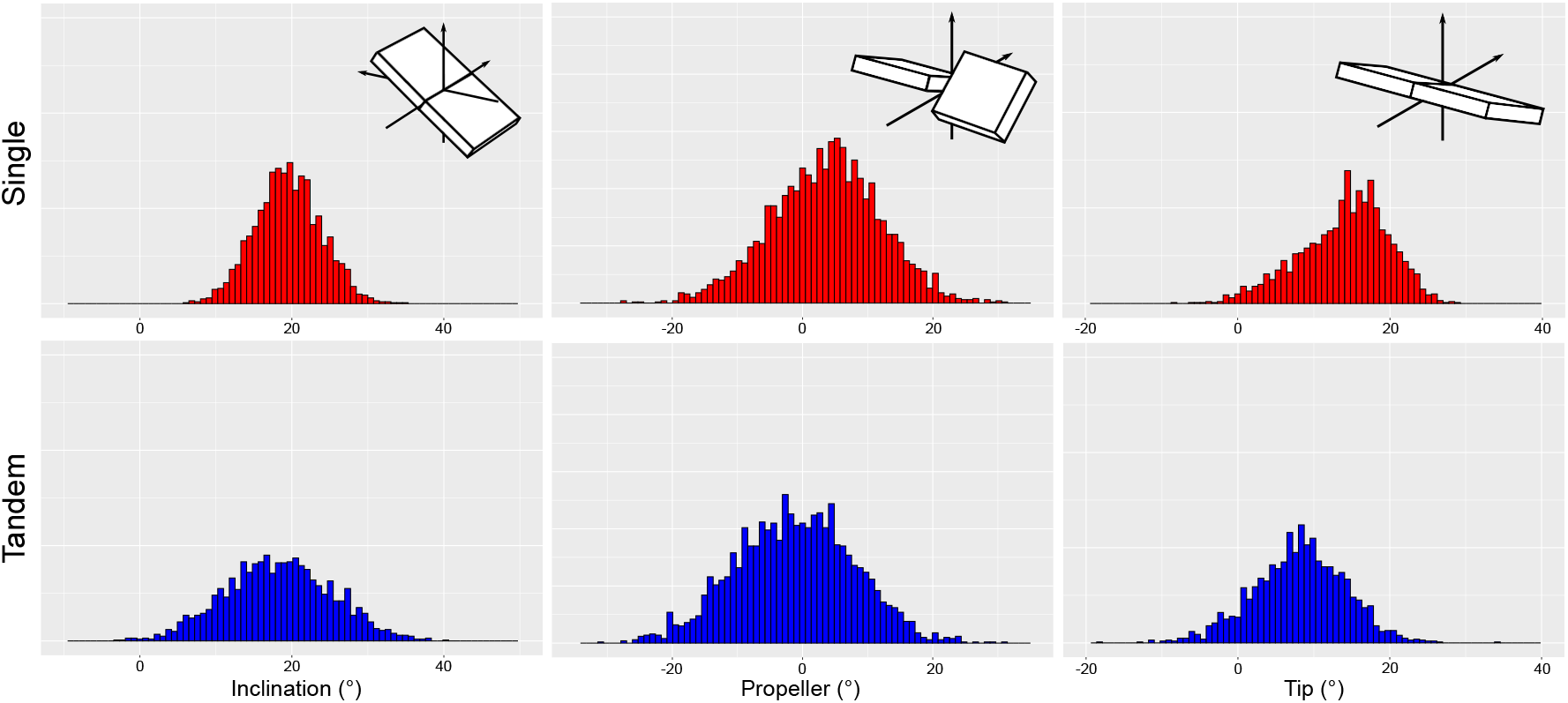
Distribution of three characteristic DNA helix intra base-pair parameters for dC8-OG19 over 2 *μ*s MD simulation, for a single 8-oxoguanine (Single, red, top) and both 8-oxoguanine and Ap site (Tandem, blue, bottom). The structural deformation with respect to canonical B-DNA is globally shier for the tandem-damaged than the singly-damaged complex.

Noteworthy, Qi *et al*^21^ reported a change in puckering values upon oxidation of a guanine residue. 8-oxoG would exhibit a C4’-exo puckering while a canonical dG ribose moiety would harbor a C2’-endo conformation. This would promote the recognition of 8-oxoG by MutM. In our simulations, the frequency of the C2’-endo conformation of OG19 is increased by the presence of tandem lesion compared to a single 8-oxoG (42.4% and 21.3%, respectively). However, rather than the C4’-exo puckering (2.9% and 7.6% for tandem- and single-damaged), the C1’-exo is the main or second preponderant conformation (42.5% and 61.5%) - see Figure S6. Concerning the inter base-pair parameters, DNA structural values are comparable for single- and tandem-damaged systems and in agreement with previous works^16^. As could be expected though, the absence of the nucleobase at position 20 upon mutation to Ap site influences the stability of the canonical stacking that is usually conserved in the singly-damaged complex. It is reflected in the distribution of the parameters values, which is much broader in the presence of Ap at position 20 - see Figure S7. This highlights the blurrier structural signature exhibited by the tandem-damaged DNA helix, which is another criteria that might affect the interaction with the surrounding amino acids, hence the efficiency of the 8-oxoG extrusion by MutM.

## Discussion

MutM, the bacterial analog of the human Fpg, is responsible for the recognition and repair of the utmost common 8-oxoG lesion. The Fpg(MutM):DNA interface has been investigated by NMR, X-ray and molecular dynamics simulations, probing the key residues that play a crucial role in the most specific recognition of 8-oxoG, but also of other DNA lesions^15,17,19–22,28,34–37^. An intercalation triad (M77, R112, F114) has been characterized, and several other residues are known to be essential in MutM:DNA interactions and 8-oxoG extrusion, guiding the lesion towards the N-terminal proline responsible for the Schiff base formation. Intrahelical insertion of a single F114 wedge residue^22,36,37^ is marked and allows a slow scanning of the double helix by MutM and analog enzymes. Among MutM key-residues, R264, located in the Zn-finger domain, is highly conserved and important for 8-oxoG extrusion^20,21^. N174 also plays a key-role and its mutation leads to the perturbation of the R264 contacts^17,22^. Besides, the C-terminal flexible loop is known to be essential for the 8-oxoG recognition by folding over the lesion in a capping process^19,21^.

While the recognition and repair of single 8-oxoG by MutM are well documented, their perturbation upon the presence of tandem lesions is very poorly understood. However, it has been shown that ionizing radiations can lead to the formation of tandem lesions^38^, rendering 8-oxoG refractory to excision by glycosylases^8,39^. Such multiple damaged sites are highly mutagenic and increase the risks of cancer development^40,41^. They can also be cytotoxic as their error-prone repair can result in the formation of deleterious double-strand breaks^42,43^. Noteworthy, the high toxicity of the DNA lesions induced by ionizing radiation is also exploited for the development of cancer (radio)-therapies^44^. In this context, we investigated the structural impact of tandem lesions on the interactions between MutM and a 13-bp oligonucleotide harboring the 8-oxoG lesion at position 19. Using molecular modeling and machine-learning analysis, we highlighted a structural re-organization of MutM canonical interaction network around 8-oxoG upon the presence of an adjacent abasic site at position 20.

The interaction network involving the intercalation triad and the damage is perturbed by the dG20 → Ap20 mutation. The MutM:DNA interactions are more pronounced, leading to a more rigid system, which could explain the difficulty of MutM to process such multiple damaged sites. While in the simulation of the singly-damaged system, the classical interaction patterns are observed, the presence of an additional Ap site results in the rotation of R76 that provokes a shift of the intercalation triad. Noteworthy, as R76 is poorly conserved in MutM sequences from different organisms^20,29^, one cannot rule out the possibility of a different reorganization around the intercalation triad. First observations of our MD trajectories allowed to describe the re-shaping of the MutM:DNA interaction patterns - see Figure 2 and 3. The structural analysis of the DNA oligonucleotide also reveals changes in the local conformation of the lesion site (see Figure 5 and Table 1), which might jeopardize the efficiency of 8-oxoG recognition by the enzyme.

In order to go beyond the visual observation of MutM:DNA interactions, we applied machine learning (ML) techniques to provide an extensive map of these contacts. ML methods have gained enormous amount of attention in recent years. Their power in finding important information out of large amount of data has been exploited by the biochemistry community, many interesting applications have been showcased in the literature. Recently, Fleetwood et al.^30^ have demonstrated its capability in learning ensemble properties from molecular simulations and providing easily interpretable metrics describing important structural or chemical features. The machine-learning analysis of our trajectories is based on the demystifying package from Fleetwood et al.^30^. Residues highlighted as providing a significant contribution to the MutM:DNA bonding by the MLP analysis are in agreement with data from the literature. Comparison of the residues importance in MutM:DNA interactions upon single or tandem lesions allowed to pinpoint the changes in the interaction patterns, which concern the most important features of MutM - see Figure 4. Apparently in contradiction with common chemical sense, MLP analysis revealed that the dG20 → Ap20 mutation leads to stronger, more stable interactions between the two macromolecules. The contribution of the residues involved in DNA anchoring is almost systematically increased in the tandem-damaged system. Nuleic acids also exhibit stronger interactions with MutM in the case of tandem lesion, which overall suggests that the presence of a second damage somehow results in a more rigid complex than when an isolated 8-oxoG is present. However, the global rigidity of the tandem-damaged MutM:DNA complex can actually be counterproductive for repair since it has been evidenced that flexibility of the DNA strand is a key feature correlating with 8-oxoG removal^29^. This consideration is also further reinforced by the fact that conversely, the catalytic N-terminal residues are less involved in the MutM:DNA complex stability in the case of tandem-damaged nucleotides. This is also the case for the 211-234 loop region which is known to play a key-role in 8-oxoG extrusion. Hence, the presence of the Ap site alongside the 8-oxoG lesion impacts the canonical structural behavior of these two important MutM regions, which might also contribute to the lower repair efficiency.

Our study provides an example of the predictive power of all-atom, MD simulations coupled to machine learning analysis, applied to a very challenging test-case. Indeed, the combination of oxidatively-generated DNA lesions embrace a combinatorial chemistry, with contrasted structural, mechanical and dynamic properties. Additionally, MutM/Fpg are very flexible proteins^29^, certainly difficult to properly sample. The efficiency of our protocols gives perspectives for its extension towards other tandem systems and the investigation of sequence effects^12,45,46^. Furthermore, the biological significance of rationalizing this complex scenario is also unquestionable. Indeed, ionizing radiations can be satisfactorily exploited in cancer therapy, and the inhibition of repair enzyme by combined chemotherapy can prove a most valuable synergy in assuring the accumulation of lesions necessary to reach the apoptosis threshold. Understanding of the molecular mechanisms underlying DNA repair is thus crucial for also offering novel perspectives for cancer research.

## Materials and Methods

### All-atom molecular dynamics simulations

All MD simulations were performed with the Amber and Ambertools 2018 packages^47^. The starting X-ray structure of *Bacillus stearothermophilus* MutM was taken from the structure obtained by Verdine and coworkers^21^, PDB ID code 3GO8. The crystallographic self-complementary ds-DNA is a 13-bp sequence d(GTAGATCCGACG).

(CGTCC**G**GATCT) featuring 8-oxoG as the 19th nucleobase (in bold). It should be noted that the *β*F-*α*10 loop 217–237 of MutM, absent from the crystal structure, was reconstructed using Modeller. The zinc atom present in the zinc-finger motif of MutM was kept and described with parameters taken from the Zinc AMBER Force Field (ZAFF) developed by Merz and coworkers^48^. 19 potassium ions were added to neutralize the MutM:DNA complex, which was embedded in a 92×97×91 Å^3^ TIP3P water molecules bath. The Amber ff14SB^49^ was used throughout, including the bsc1 force field corrections for the DNA duplex^50^.

The parameters for 8-oxoG and Ap site have been generated with a standard antechamber procedure embedded in Amber 18^47^, as described in previous references^51–53^ and in agreement to the literature. Four 10,000 steps minimization runs were carried out on the initial MutM:DNA complex, imposing restraints on the amino and nucleic acids, that were gradually decreased from 20 to 0 kcal/mol/Å^2^ along the four runs. The temperature was then raised from 0 to 300K in a 50 ps thermalization step, and afterwards kept constant using the Langevin thermostat with a collision frequency *γ*ln of 1 ps^*−*1^. The system was subjected to a 1 ns equilibration run in the NPT ensemble. Finally, two replica of 1 *μ*s production run were performed to sample the conformational ensemble of the system. The Particle Mesh Ewald method was used to treat electrostatic interactions, with a 9.0 Å cutoff.

The structural descriptors of the DNA helix were evaluated based on a post-processing analysis with Curves+^32^ and other distance and RMSD values were monitored using Ambertools.

### Multilayer Perceptrons analysis

The Multilayer Perceptrons (MLP) is a fully connected artificial neural network (ANN) with one input layer, one output layer and at least one hidden layer. After tests, the architecture of the MLP was chosen to contain a single layer of 200 neurons to provide good accuracy. The rectified linear unit function (ReLU)^54^ was used for the activation of neurons, and the Adam algorithm was used for optimization. The inverse of the distances between the geometric centers of the residues were used as the input features for the multilayer perceptrons neural network, due to better overall performance over Cartesian coordinates, according to Fleetwood et al^30^. These internal coordinates were computed for all residue pairs and all frames. Each frame of the trajectories was labelled as either 1 or 0 according to whether the distance between the DNA lesion(s) and the protein is lower (bounded) or higher (non-bounded) than 10 Å. These sets of input features and labels were fed to the MLP classifier for training. Upon completion of the training, layerwise relevance propagation (LRP) was performed to find out the important features of the DNA/MutM interface.

## Acknowlegements

Support from ENS de Lyon is gratefully acknowledged. This work was performed within the framework of the LABEX PRIMES (ANR-11-LABX-0063) of Université de Lyon, within the program “Investissements d’Avenir” (ANR-11-IDEX-0007) operated by the French National Research Agency (ANR).

## Notes

### Competing Interest Statement

The authors have declared no competing interest.

