## Supporting Materials for "Recognition of a Tandem Lesion by DNA Glycosylases Explored Combining Molecular Dynamics and Machine Learning"

### Supporting Materials for: Recognition of a tandem lesion by DNA glycosylases explored through Molecular Dynamics simulations and a Machine Learning protocol

Emmanuelle Bignon,[*,†] Natacha Gillet,[†] Chen-Hui Chan,[†] Tao Jiang,[†] Antonio Monari,[‡] and Elise Dumont[†,¶]

†*Univ Lyon, ENS de Lyon, CNRS UMR 5182, Université Claude Bernard Lyon 1, Laboratoire de Chimie, F69342, Lyon, France*

‡*Université de Lorraine and CNRS, LPCT UMR 7019, 54000 Nancy, France*

¶*Institut Universitaire de France, 5 rue Descartes, 75005 Paris, France*

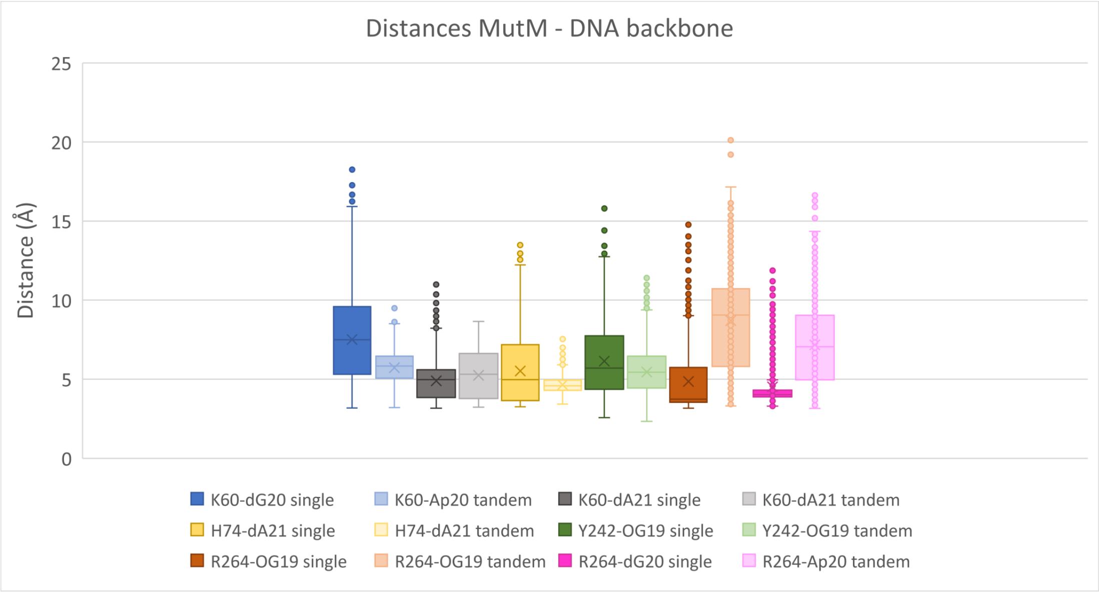


Figure S1 – Distribution of the distances between DNA phosphate groups and MutM amino acids stabilizing the MutM:DNA complex, for singly- (dark colors) and tandem- (light colors) damaged systems.

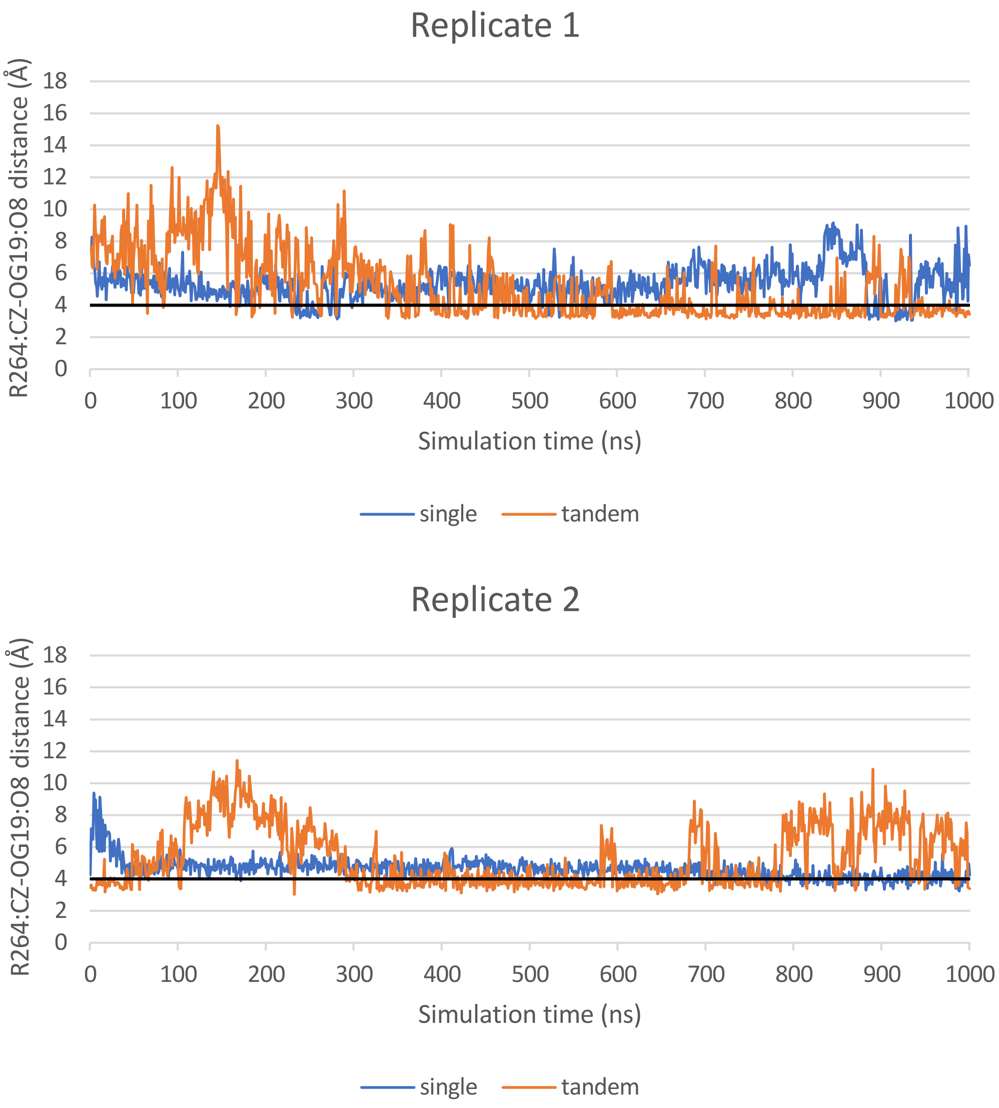


Figure S2 – Evolution of the OG19:O8-R264:CZ distance along the two 1$\mu$s MD replicates. Singly- and tandem-damaged systems are depicted in blue and range, respectively. The threshold value for the H-bond to be effective is set to 4Å by observation of the trajectories.

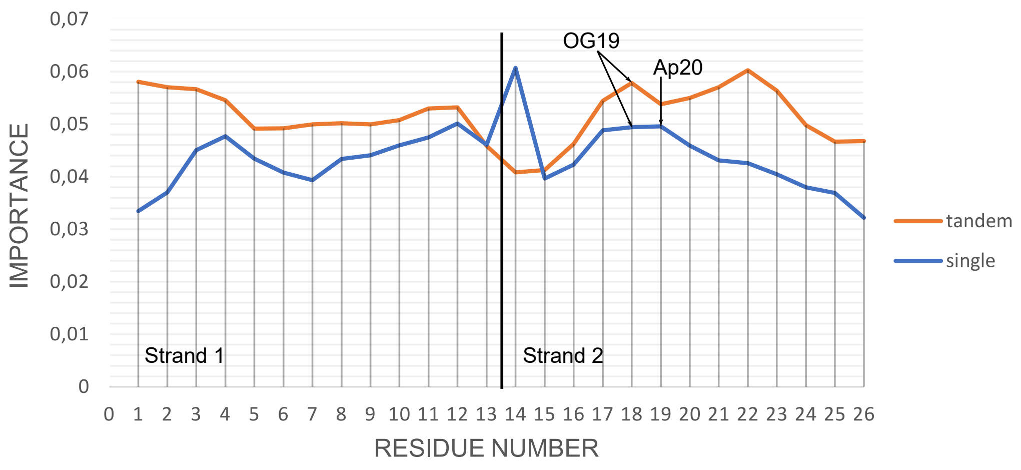


Figure S3 – Importance values of the DNA helix nucleotides denoting their contribution to the MutM:DNA bonding, for singly- (blue) and tandem- (orange) damaged systems. The vertical line separates strand 1 (residues 1 to 13) from strand 2 (residues 14 to 26), and the lesions sites (OG19 and Ap20) are pinpointed by black arrows.

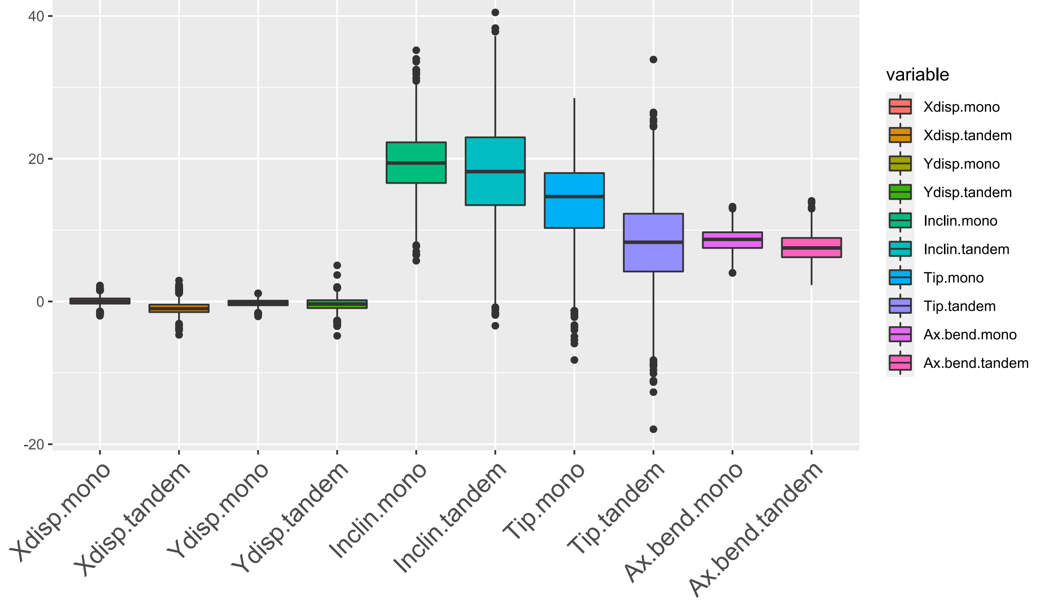


Figure S4 – Backbone parameters for dC8-OG19 for systems harboring either a single 8-oxoG (single) or 8-oxoG + Ap (tandem).

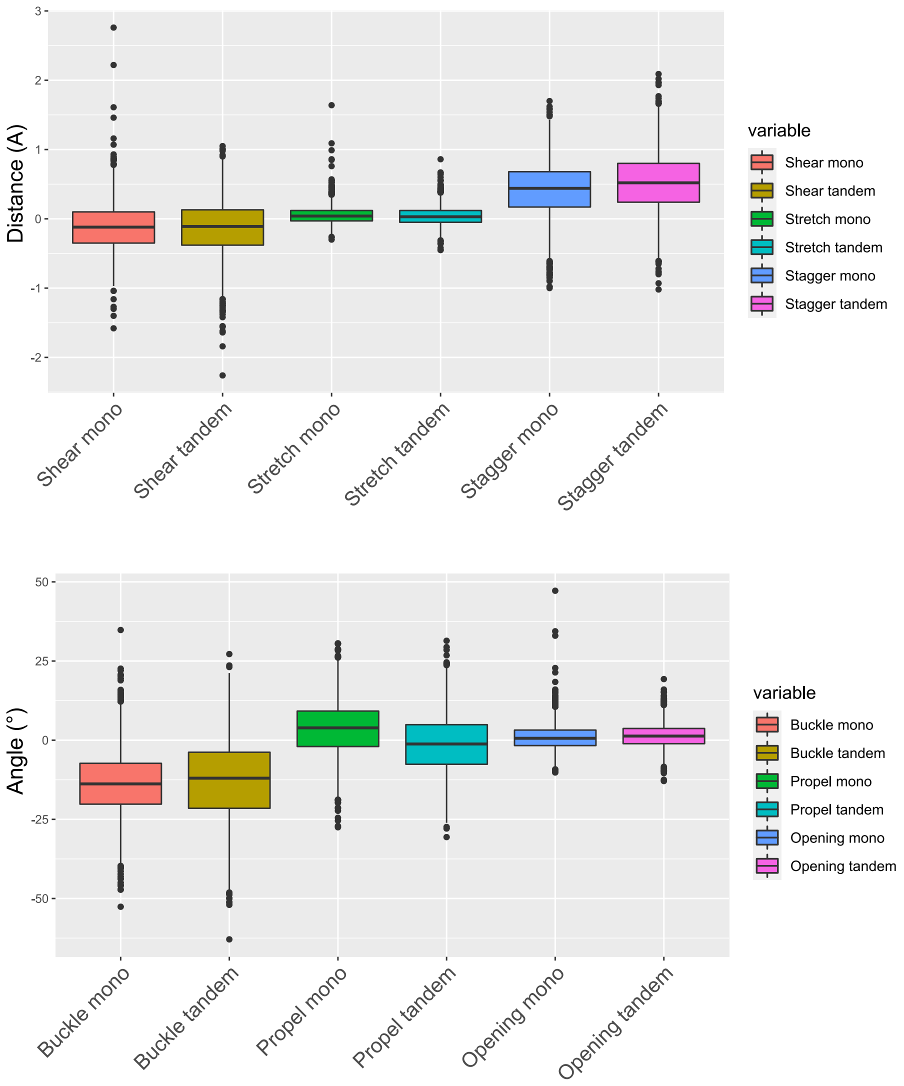


Figure S5 – Intra base-pair parameters for the dC8-OG19 base-pair for systems harboring either a single 8-oxoG (single) or 8-oxoG + Ap (tandem).

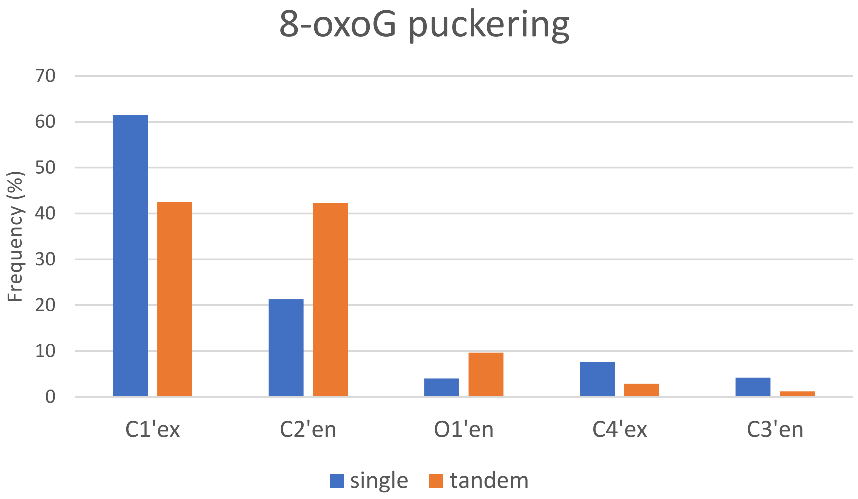


Figure S6 – Frequency of OG19 puckering over the MD simulations for the singly- (blue) and tandem- (orange) damaged systems.

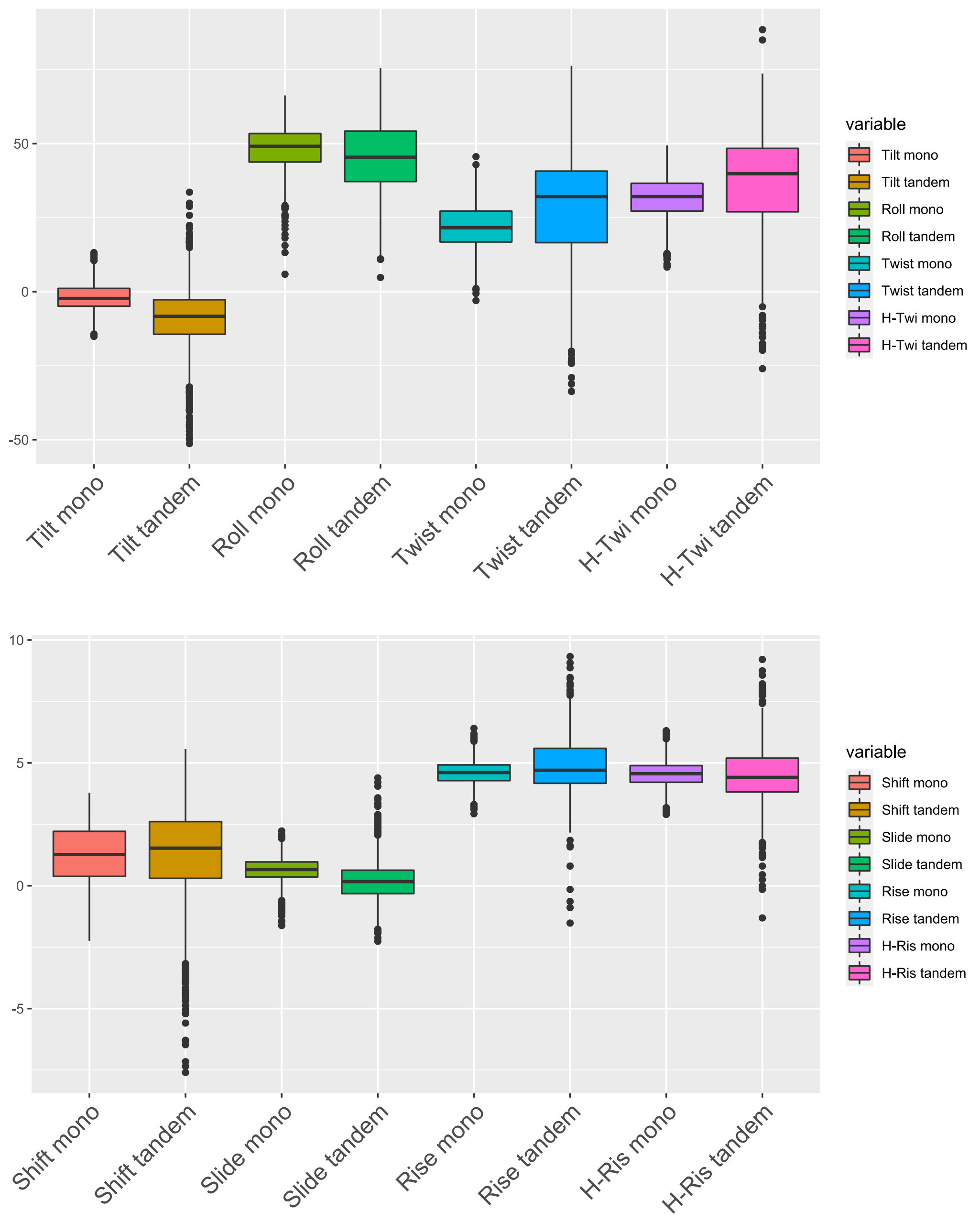


Figure S7 – Inter base-pair parameters for dC7-dG20 / dC8-OG19 for systems harboring either a single 8-oxoG (single) or 8-oxoG + Ap (tandem)
